# WDR23 mediates NRF2 proteostasis and cytoprotective capacity in the hippocampus

**DOI:** 10.1101/2023.10.10.561805

**Authors:** Jiahui Liu, Chatrawee Duangjan, Ronald W. Irwin, Sean P. Curran

## Abstract

Pathogenic brain aging and neurodegenerative diseases such as Alzheimer’s disease and Parkinson’s disease are characterized by chronic neuroinflammation and the accumulation of dysfunctional or misfolded proteins that lead to progressive neuronal cell death. Here we demonstrate that a murine model with global loss of the CUL4-DDB1 substrate receptor WDR23 (*Wdr23KO*) results in changes in multiple age-related hippocampal-dependent behaviors. The behavioral differences observed in *Wdr23KO* animals accompany the stabilization of the NRF2/NFE2L2 protein, an increase in RNA transcripts regulated by this cytoprotective transcription factor, and an increase in the steady state level of antioxidant defense proteins. Taken together, these findings reveal a role for WDR23-proteostasis in mediating cytoprotective capacity in the hippocampus and reveal the potential for targeting WDR23-NRF2 signaling interactions for development of therapies for neurodegenerative disorders.

**HIGHLIGHTS:**

- WDR23 regulates NRF2/NFE2L2 stability in the mouse hippocampus
- Loss of *Wdr23* significantly increases the expression of NFE2L2/NRF2 target genes
- Global loss of WDR23 influences age-related behaviors differentially in males and females

## INTRODUCTION

The NFE2L2/NRF2 transcription factor functions as a regulator of oxidative stress and neuroinflammation in addition to many other metabolic responses [1–3]. Although a large body of evidence documents the proteostatic regulation of NRF2 by CUL3-KEAP1, regulation by the CUL4-DDB1-WDR23 E3-ubiquitin ligase also plays important roles [4, 5]. NRF2 is an important homeostat that manages oxidative stress resistance pathways that are critical for health across tissues and throughout the lifespan.

Oxidative stress, a result of the imbalance between free radicals and antioxidants, is recognized as a fundamental contributor to the aging process [6]. The brain exhibits a pronounced requirement for ATP originating from mitochondrial metabolism, resulting in constant generation of free radicals and reactive oxygen species (ROS) [7]. Neurons, in particular, exhibit a heightened susceptibility to oxidative stress-induced damage over the lifespan and as such, sophisticated are utilized to alleviate the oxidative-stress condition [8]. Accumulating evidence has shown that excessive oxidative stress can lead to the gradual deterioration of brain tissues, which contributes to age-related cognitive decline and an increased vulnerability to many disorders [9, 10].

In this context, NRF2 emerges as a central player in the aging narrative. NRF2 acts as a transcription factor that regulates the expression of antioxidant and detoxification genes, thereby helping to mitigate oxidative stress and inflammation [1, 11–13]. Beyond its role in oxidative stress resistance, NRF2 has been increasingly recognized for its involvement in age-related diseases such as Alzheimer’s disease (AD), Parkinson’s disease (PD), and Huntington’s disease (HD) [14–16]. Thus, understanding the mechanisms through which NRF2 and other oxidative stress resistance pathways modulate brain aging, is paramount in advancing our knowledge of neurodegenerative disorders and promoting healthy aging.

An emerging body of evidence has shed light on the discrepancies in redox signaling between the sexes [17, 18]. These sex-related differences in coping with oxidative stress extend beyond normal physiological context, and confer distinct advantages to females, including a longer lifespan and reduced susceptibility to many diseases characterized by oxidative stress [17, 19, 20]. However, it is worth highlighting an augmented vulnerability to AD among females [21, 22]. Research indicates that a number of factors, including sex hormones [23–25], sex chromosome genes [26, 27], mitochondrial functions [28, 29] and their interplays can impact brain function and cognitive aging differently compared to males. Nevertheless, the precise mechanistic underpinnings as to why females are protected from certain oxidative stress-related diseases but vulnerable toothers, remains an unexplored area of research.

In this study, we examine the impact of genetic disruption of *Wdr23* on age-related changes in behavior in the murine model, and document changes in the NRF2 homeostat in the hippocampus. Collectively, this work reveals a conserved role for WDR23-dependent proteostatic control of NRF2 in the nervous system, which provides a new model for manipulating NRF2 activity in the aging brain that can complement models disrupting KEAP1 regulation and result pleiotropic outcomes.

## MATERIALS AND METHODS

### Animals

All animal protocols were approved by the Institutional Animal Care and Use Committee (IACUC) of the University of Southern California and all the procedures were conducted in compliance with institutional guidelines and protocols.

*Wdr23* knock-out (*Wdr23KO*) mice were generated by Wellcome Trust Sanger Institute [30, 31]. *Wdr23KO* animals were subsequently backcrossed nine times into our C57BL/6J (WT) strain from the Jackson laboratory. Heterozygous (*Wdr23 +/−*) *dams* and *sires* were then mated to generate *Wdr23+/*+ and *Wdr23−/−* animals that were maintained as WT and KO, respectively. Mice were kept in a 12:12 h light-dark cycle, constant temperature, and humidity room. All animals were allowed *ad libitum* access to water and food.

### Rodent diets

Mice were fed ad lib with irradiated D12450K rodent chow (Research Diets), containing 16.1kJ/g of digestible energy (animal-based protein 3.22kJ/g, carbohydrate 11.27kJ/g, fat 1.61 kJ/g).

### RNA extraction and RNA sequencing

The hippocampus tissue of 44 weeks old male mice was collected and lysed in RNeasy Lipid Tissue mini kit reagent (QIAGEN). RNA was extracted according to the manufacturer’s protocol. Qubit RNA BR Assay Kit was used to determine RNA concentration. Isolated RNA was sent to Novogene for library preparation and deep sequencing in biological replicates (N=6 WT, N=4 *Wdr23KO*). The read counts were then used for differential expression (DE) analysis by using the R package DEseq2 (R version 3.5.2). Differentiated expressed genes were analyzed using p value <0.05 and fold change >1.5 as cutoff.

### Western blot analysis

The hippocampus tissue of 8-14 weeks old male mice was collected for WB analysis. Whole cell lysates were prepared in M-PER buffer (1x Mammalian Protein Extraction Reagent (Thermo Scientific), 0.1% Halt Protease & Phosphatase inhibitor (Thermo Scientific) according to the manufacturer’s protocol. Total protein concentrations were quantified by Bradford assay (Sigma). An equal amount of protein (20 µg) was separated on 4%-12% bis-tris polyacrylamide gel (Invitrogen,) in MOPS running buffer (Invitrogen) and then transferred to nitrocellulose membranes (GE Healthcare Life science). After blocking for 1 h with 3% BSA in PBST (PBS, 0.1% Tween 20), the membranes were subjected to immunoblot analysis. Antibodies used include: NFE2L2/NRF2 (Proteintech, 1:1000), ALDH2 (Proteintech, 1:1000) GCLM (Proteintech, 1:1000), NQO1 (Proteintech, 1:1000), GPx4 (Proteintech, 1:1000), GPx1 (Proteintech, 1:1000)), GSTA4 (Proteintech, 1:500), SOD1 (Proteintech, 1:1000), SOD2 (Cell signaling, 1:1000), G6PD (Proteintech, 1:1000), β-actin (Millipore Sigma, 1:10000) and HRP-conjugated secondary antibodies (Thermo Fisher, 1:10,000). Specific protein bands were visualized and evaluated using FluorChem HD2 (ProteinSimple).

### Behavioral Assays

Prior to behavioral tests, mice were habituated to the testing room for 30 minutes. Arenas and objects were thoroughly cleaned between mice using 70% (vol/vol) ethanol. Test trials were video recorded and analyzed using Noldus Ethovision XT software (Version 16).

#### Spontaneous Alteration Tests (SAB)

The arena consisted of three arms (x cm) spaced at 120-degree angles. Mice were placed in one of the arms and allowed to explore freely for 5 minutes. An arm entry was considered when the animal placed all four paws into the arm. Three consecutive entries into different arms were considered an alteration. The total number of arm entries was recorded, and the percentage of correct alterations was calculated.

#### Open Field Tests

The open field tests were conducted the day after SAB. Mice were gently placed in the center of a square arena (40 × 40 cm) and allowed to explore freely for a duration of 10 minutes. The arena was divided into 16 equal squares, with the center area defined by the middle 4 squares. Throughout the trial, total distance moved, the overall time the mice remained mobile, and the time spent in the center area were recorded. To calculate the average velocity, the total distance moved was divided by the total time of mobility.

#### Novel object placement and recognition

We employed a four-day protocol to evaluate the recognition memory. Same arena in the open field test was used. Open field test was considered as day 1 habituation of novel object placement and recognition tests. On day 2, each animal was given another habituation trial performed as day 1. On day 3 (familiarization and novel object placement), the test arena was prepared with two identical objects positioned in adjacent corners. The mice were placed in the arena, facing opposite to the objects, and allowed to explore for 10 minutes. After 1 hour, the animals were reintroduced to the arena, but with one of the identical objects moved to the diagonally opposite corner. The mice were again allowed to explore for 10 minutes. On day 4 (novel object recognition), 24 hours after the familiarization trial, mice were placed in the arena and allowed to explore for 10 minutes with one of the familiar objects replaced with a novel object. Throughout all trials, exploration time for each object was recorded. Percentage of time exploring each object was calculated.

### Physiological cages

Metabolic assessments were conducted using the PhenoMaster system (TSE-Systems) as previously described [32]. One week after the last behavioral test, mice were individually housed in PhenoMaster cages and subjected to automated monitoring over a span of four days. The initial day was designated as an acclimation period, and data were computed as the mean values obtained from days 2 to 4. Ambulatory activity was calculated as total counts of breaking beams in the x, y, and z axes. Respiratory exchange ratio (RER) was calculated as the ratio between production of carbon dioxide and consumed volume of oxygen. Energy expenditure in each animal was normalized by their whole-body weight.

### Primary cell culture

Primary cortical neurons or mixed glia were isolated from embryonic day 17 (E17) fetuses from timed-pregnant C57BL/6J WT or *Wdr23KO* age-matched mice. Briefly, brains were removed in cold Hanks Balanced Salt Solution HBSS containing 10 mM HEPES, cerebral cortices including hippocampus were dissected from 5-8 brains, meninges removed, and incubated for 10 min at 37°C in trypsin containing HBSS. Cells were dissociated with pipet then poured through a 40-micron filter. Cells were counted by hemacytometer then seeded at a density of 1 million cells per well in 6-well poly-D-lysine pre-coated plates (Greiner Bio-One) and incubated at 37°C, 5% CO_2_. Neurons were cultured for 7-10 days in complete Neurobasal Plus Medium (Gibco). Mixed glial cells were cultured with the same cortical cell isolation procedure, first seeded in T75 flasks (Corning) and grown for approximately 2 weeks to reach 80-90% confluency in Dulbecco’s Modified Eagle Medium (DMEM; Gibco) with 10% fetal bovine serum and 1% antibiotic-antimycotic (Corning). Mixed glial cultures adhered to T75 flask were then trypsinized and re-plated in 6-well plates at a density of 1 million cells per well and observed to be 80-90% confluent. Cells were harvested 24 hours after fresh media changes. Cells were rinsed in warm Dulbecco’s Phosphate Buffered Saline (DPBS) to remove media and debris then lysed with M-PER Mammalian Protein Extract Reagent (Thermo) containing Halt protease inhibitors (Thermo). Total cell lysates were collected on ice followed by centrifugation (10,000 x g for 10 min at 4°C) to extract soluble proteins for immunoblots.

### Statistical Analysis

All experiments were performed at least in triplicate. Data are presented as mean ± SEM. Data handling and statistical processing were performed using GraphPad Prism 8.0. In SAB and open field test, data were analyzed using two-way ANOVAs with genotype and age as between-subjects factors. For novel object placement and recognition, percentage of time exploring moved or novel object in each group was compared with 50% using one-sample t-test.

Measurements from physiological cages were analyzed using either repeated two-way ANOVAs across time points and genotype as a between-subjects factors, or two-way ANOVAs with genotype and age as between-subjects factors. All other comparisons between two groups were done using unpaired Student’s t-test. Differences were considered significant at the p ≤ 0.05 level. *p<0.05, **p<0.01, ***p<0.001, ****p<0.0001, compared to C57BL/6J (WT) control or indicated with brackets.

## RESULTS AND DISCUSSION

### *Wdr23KO* animals display age-related differences in behavior

A decline of neuronal proteostasis during natural aging or extreme stress can lead to neuronal disorders [33]. To investigate the impact of the proteostasis regulator *Wdr23* on aging-related changes in behavior, we conducted a series of cognitive assessments in a mouse model lacking *Wdr23* (*Wdr23KO*) and compared them with wild-type (WT) controls at different ages (young, 4-month-old; middle/late-aged, 16-month-old).

Firstly, we employed the novel object recognition (NOR) and placement (NOP) tasks (**Figure 1A**) to gauge cognitive function in these animals (**Figure 1** and Figure S1). The NOR task assesses the ability to distinguish object identity, a cognitive function reliant on various brain regions [34]. In our observations, we noted an enhanced cognitive recognition memory capacity in both old male (**Figure 1B**) and female (**Figure 1C**) animals. In the NOP task, which evaluates spatial recognition and the capacity to detect changes in the arrangement of objects [35], young *Wdr23KO* of both sexes initially displayed limited discrimination ability regarding the newly positioned object in the NOP task. However, it is noteworthy that this capacity became more pronounced in older *Wdr23KO*. In contrast, WT animals showed an age-related decline of performance in this task, as only young WT animals can discriminate the newly positioned object (**Figure 1D-E**). These findings indicate that the impact of *Wdr23* deficiency on cognitive function varies depending on the specific cognitive domain assessed, emphasizing the complexity of cognitive processes and their modulation by *Wdr23* during aging.

**Figure 1.**
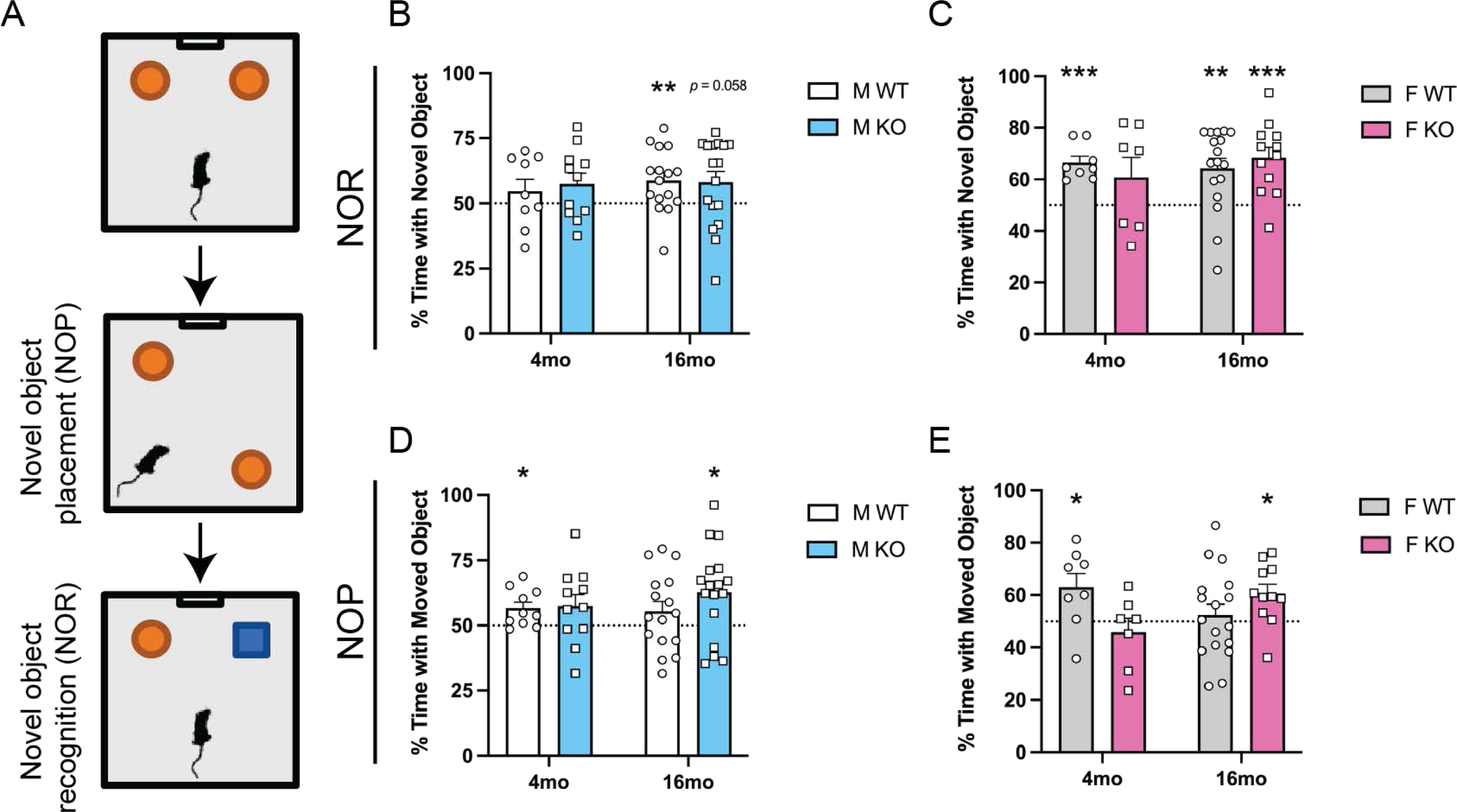
*Wdr23* effects on recognition memory. **(A)** Schematic representation of the experimental novel object placement (NOP) and recognition tests (NOR) experimental designs. Percentage of time exploring the novel object in **(B-C)** NOR and moved object in **(D-E)** NOP tests in (**B,D)** male and **(C,E)** female mice. Data are presented as mean (+SEM) values. n=7-17/group. * denotes p < 0.05, ** denotes p < 0.01.

We next assessed spontaneous alternating behavior (SAB) as a measure of exploratory behavior and spatial working memory (**Figure 2A**). Our analysis of the SAB task revealed no statistically significant differences between male WT and *Wdr23KO* animals concerning the percentage of correct alterations in the SAB task (**Figure 2B**). However, in the case of female mice, the absence of *Wdr23* was associated with improved SAB performance (**Figure 2C**). suggesting enhanced spatial working memory. Notably, young *Wdr23KO* males exhibited elevated levels of activity in such tests, as evidenced by an increased number of arm entries (Figure S2A). Conversely, young WT females displayed a significant decline in arm entries with age, a trend that was not observed in *Wdr23KO* animals (Figure S2B). It remains possible that these trends, while not statistically significant at this stage, may potentially become significant at a more advanced age. These findings emphasize the gender-specific and age-related variations in exploratory behavior and spatial working memory in response to *Wdr23* deficiency. Further investigation and longitudinal studies may shed additional light on these trends as animals continue to age.

**Figure 2.**
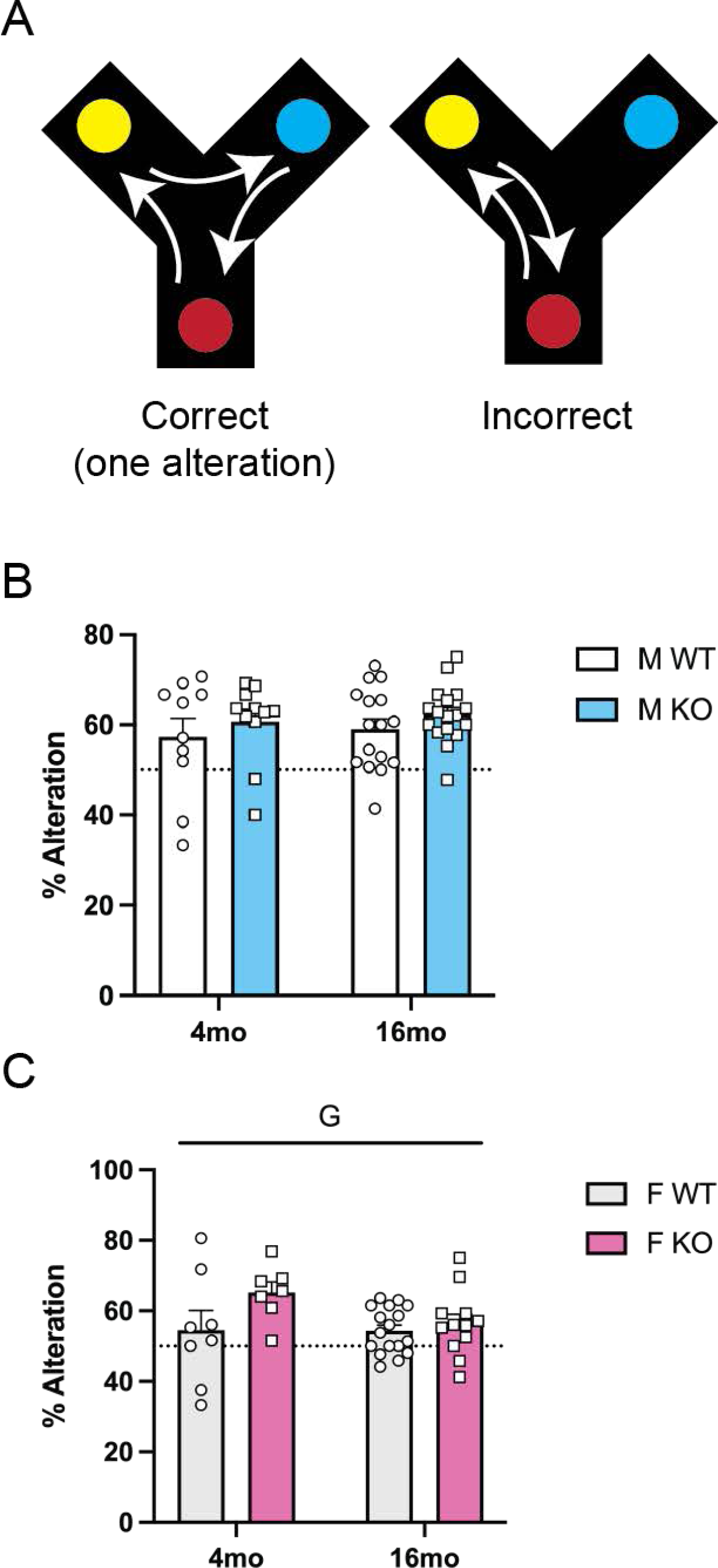
*Wdr23* effects on short-term working memory. **(A)** Illustrative cartoons showing examples of a correct and an incorrect spontaneous alternation behavior (SAB). Percentage of correct alternations were calculated in **(B)** male and **(C)** female mice. Dashed lines represent the 50% alteration level. Data are presented as mean (+SEM) values. n=7-17/group. Statistically significant main effect is denoted G (genotype).

Lastly, to gain insights into general explorative activity levels, gross locomotor activity, and exploration habits, we conducted the open field task (**Figure 3A**). In both older males (**Figure 3A-B**) and females (**Figure 3C-D**), *Wdr23KO* animals displayed notable differences in behavior. One key metric we examined was the amount of time spent in the center of the field, which is often indicative of anxiety levels in rodents [36]. Interestingly, *Wdr23KO* animals, in both sexes, spent more time in the center, suggesting a reduced state of anxiety compared to their WT counterparts. Furthermore, we assessed overall exploratory activity by measuring the total distance moved and the total time spent in motion, providing insights into locomotor activity and the propensity to explore the novel environment (Figure S3). Aged male *Wdr23KO* animals exhibited an overall reduction in exploratory activity when compared to their WT counterparts, as evidenced by decreased total distance moved and reduced total time spent in motion (Figure S3A-B). Similar to the observation in the SAB test, overall female *Wdr23KO* animals displayed diminished movement, a characteristic that did not show a further decline with age (Figure S3DF). The absence of an age-related decline might suggest that *Wdr23* does not play a role in these behaviors, or that the impact of WDR23 action does not manifest at middle age and that old animals (> 2 years) might be more informative. Nevertheless, the modest differences in the behavioral assessments reveal that loss of *Wdr23* can influence animal behaviors with age, relative to age-matched WT controls.

**Figure 3.**
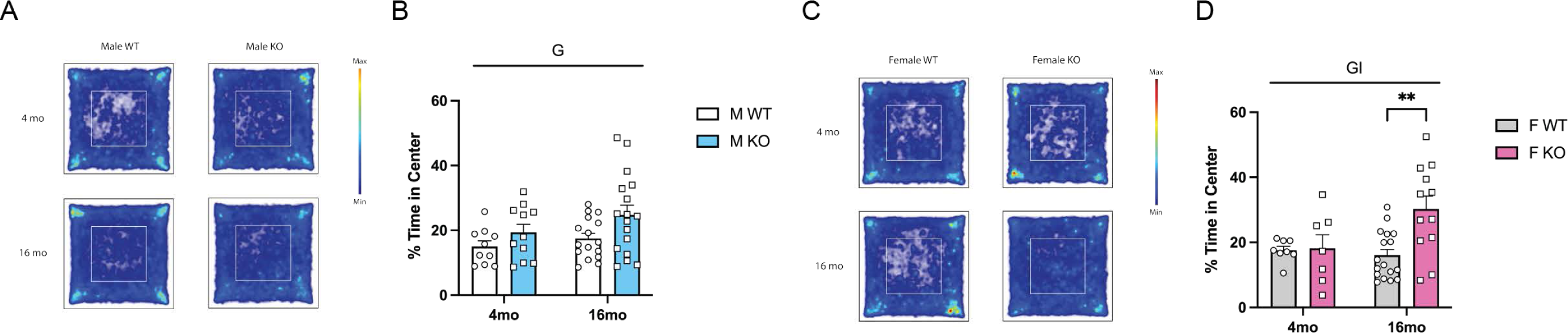
*Wdr23* effects on exploration behavior and anxiety. The heatmaps of the activity of **(A)** male and **(C)** female mice in the open field test. Percentage of time spent in the center in **(B)** male and **(D)** female mice. Data are presented as mean (+SEM) values. n=7-17/group. Statistically significant main effects are denoted by G (genotype) and I (age and genotype interaction). ** denotes p < 0.01.

### *Wdr23KO* animals have altered home-cage activity and respiratory exchange ratio

Recent findings suggest that *Nrf2* plays a role in regulating energy metabolism [37, 38]. Therefore, we accessed home-cage physical activity, respiratory exchange ratio (RER), and energy expenditure using physiological cages in *Wdr23KO* and WT animals of both sexes. In young male mice, it was evident that WT animals exhibited higher levels of spontaneous activity when compared to *Wdr23KO* mice (**Figure 4A**). However, as mice aged, we noticed an age-related increase in activity exclusively in *Wdr23KO* animals (**Figure 4C-D**). Conversely, in the case of female mice, we found that WT animals demonstrated higher activity levels than *Wdr23KO* mice, although this difference was only notable in the older groups (**Figure 4J**). Interestingly, the aging process was linked to a decrease in activity, but this reduction was specifically observed in *Wdr23KO* female mice (**Figure 4K-L**). These findings highlight the potential role of *Wdr23* in modulating physical activity levels, which, in the context of energy metabolism, can have implications for overall metabolic health and the aging process.

**Figure 4.**
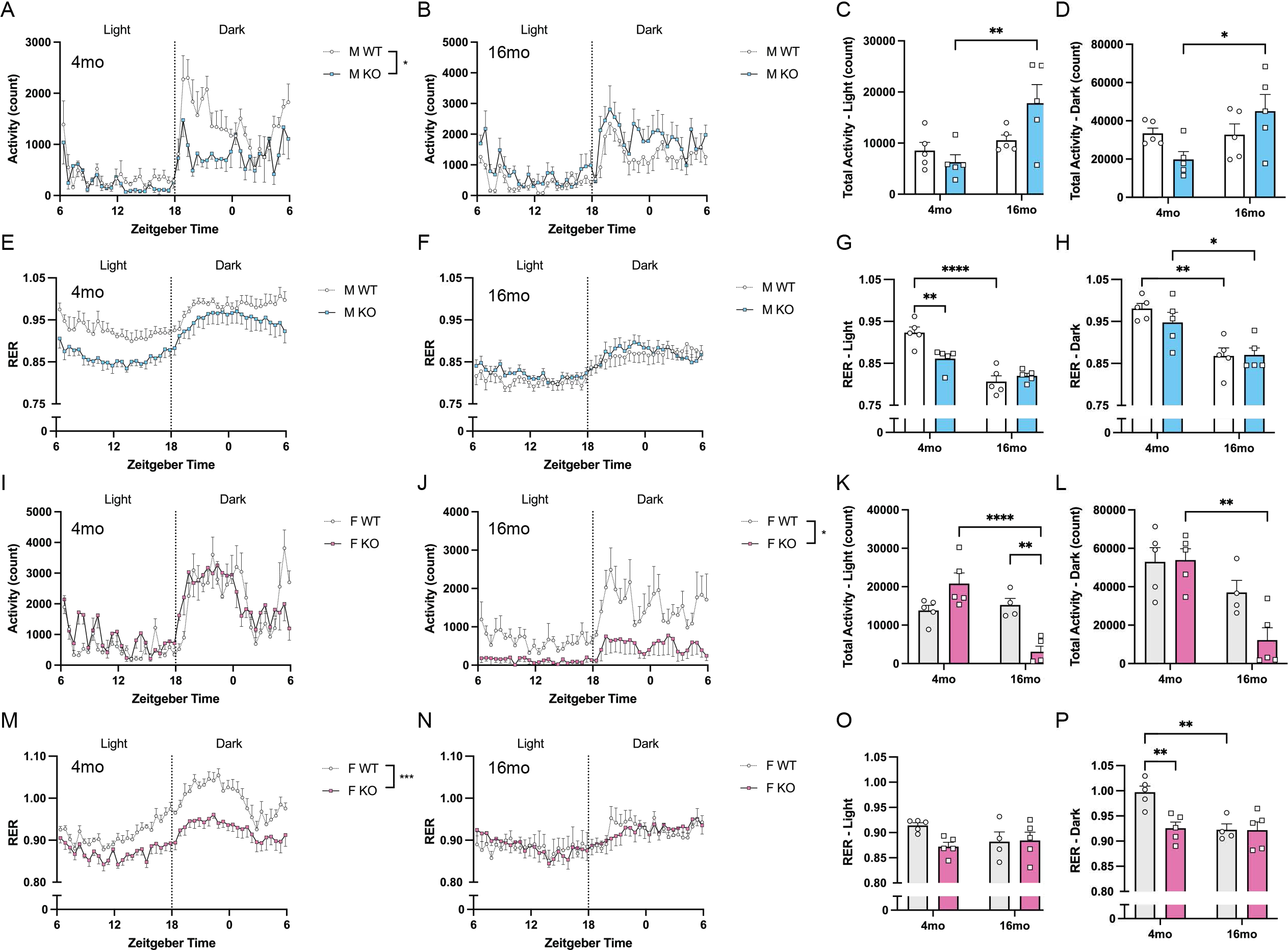
*Wdr23* effects on respiration exchange ratio and activity. Total activity and respiratory exchange ratio (RER) were measured in physiological cages. Averaged activity at different time points during the last three days in **(A)** 4-month-old male, **(B)** 16-month-old male mice. Total activity in different ages of male mice during **(C)** daytime and **(D)** nighttime. Averaged RER in **(E)** 4-month-old male, **(F)** 16-month-old male mice. Averaged RER in different ages of male mice during **(G)** daytime and **(H)** nighttime. **(I-K)** Averaged activity and **(M-P)** RER were also measured in **(I,M)** 4-month-old female and **(J-N)** 16-month-old female mice. Data are presented as mean (+SEM) values. n=4-5/group. * denotes p < 0.05, ** denotes p < 0.01, ** denotes p < 0.0001.

During the daytime, but not during nighttime, young male *Wdr23KO* animals exhibited significant lower RER when compared to aged-matched WT counterparts. Based on this difference, WT animals display an increased utilization carbohydrates as compared to *Wdr23KO* mice (**Figure 4E, G**). We also observed an age-related decline of RER in male WT animals during both daytime and nighttime, but this difference was only evident during nighttime in *Wdr23KO* animals (**Figure 4H**). Similarly, young female *Wdr23KO* animals also displayed reduced RER levels (**Figure 4M**). Notably, only female WT demonstrated a decline in RER with age during nighttime, a pattern was not observed in female *Wdr23KO* animals (**Figure 4P**). The absence of *Wdr23* had a notable impact on the change in energy expenditure with age in male mice. Specifically, old male *Wdr23KO* animals, but not WT animals, exhibited lower energy expenditure than their younger counterparts (Figure S4A-B). However, in female mice, neither age nor the presence or absence of *Wdr23* seemed to significantly affect energy expenditure (Figure S4C-D). This sex difference is of great interest and uncovering why males and females exhibit a dimorphic response to the loss of *Wdr23* could add to our understanding of sexual dimorphism in the brain and in healthy aging in general.

### *Wdr23KO* animals activate hippocampal NFE2L1/NFE2L2

The brain is under the constant need to maintain homeostasis (*e.g.*, energetic [39], proteostatic [33], redox [8]) that is mediated in part at the level of transcriptional homeostasis. To discern whether the loss of *Wdr23* resulted in a specific molecular response in the hippocampus, we performed RNAseq on dissected hippocampi from *Wdr23KO* mice and compared to samples from age-matched WT animals. 46 genes displayed a significant difference in expression (**Figure 5A**, **Table 1**) of which over 40% (20/46) have NFE2L1/NFE2L2 binding sites in the promoter region of the gene (**Figure 5B**).

**Figure 5.**
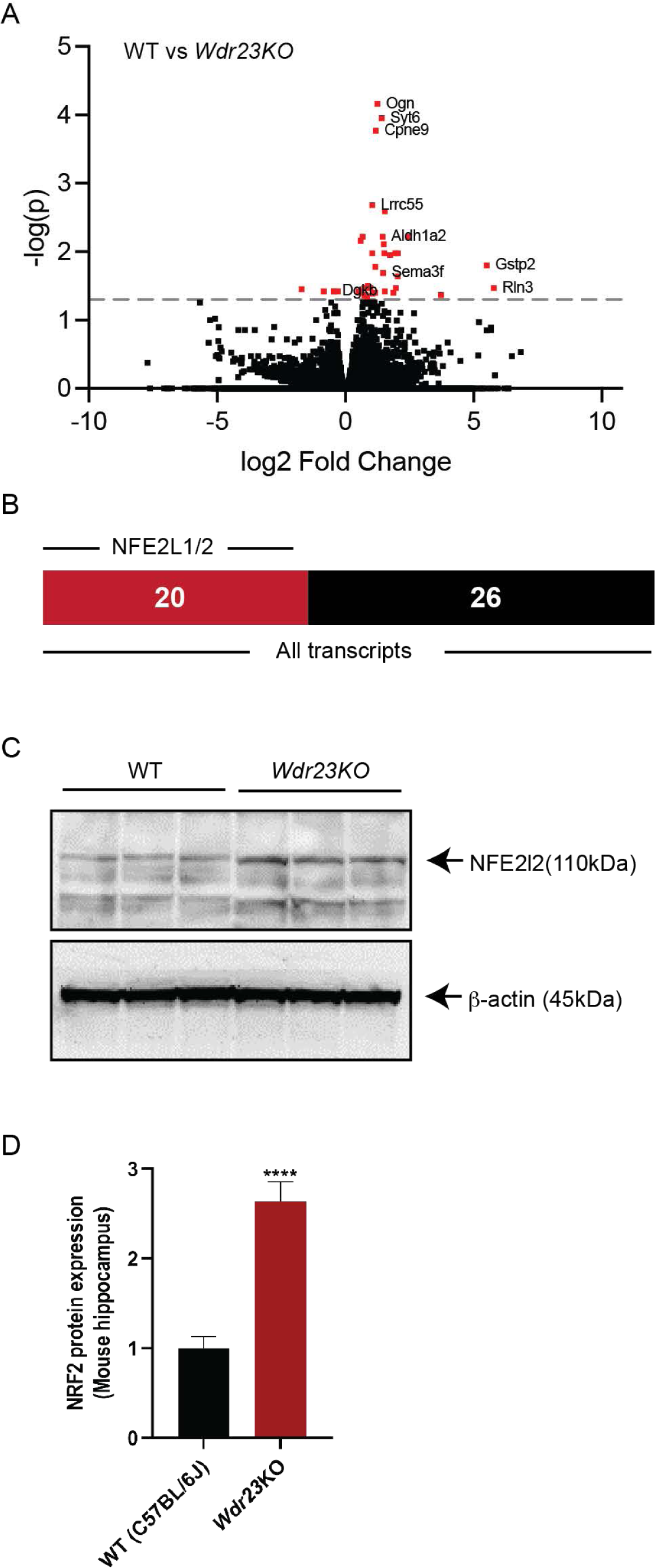
Genetic loss of *Wdr23* increases hippocampal NFE2L2/NRF2 activity. Volcano plot of differentially expressed genes between WT and *Wdr23KO* **(A)**. Each gene is represented as single dot. The mean expression level for each gene is indicated by log2FoldChange. Genes that reached the significance threshold (adjusted p-value < 0.05) are shown as red dots. Of the 46 gene transcripts with a significant difference in expression, 20 of these genes contain NFE2L1/NFE2L2 binding elements [44]. **(B)**. Western blot analysis of NFE2L2/NRF2 **(C, D)** protein revealed increased steady-state levels in *Wdr23KO* male mice hippocampus compared to age matched WT (C57BL/6J) control. ****p<.0001

**Table 1.**
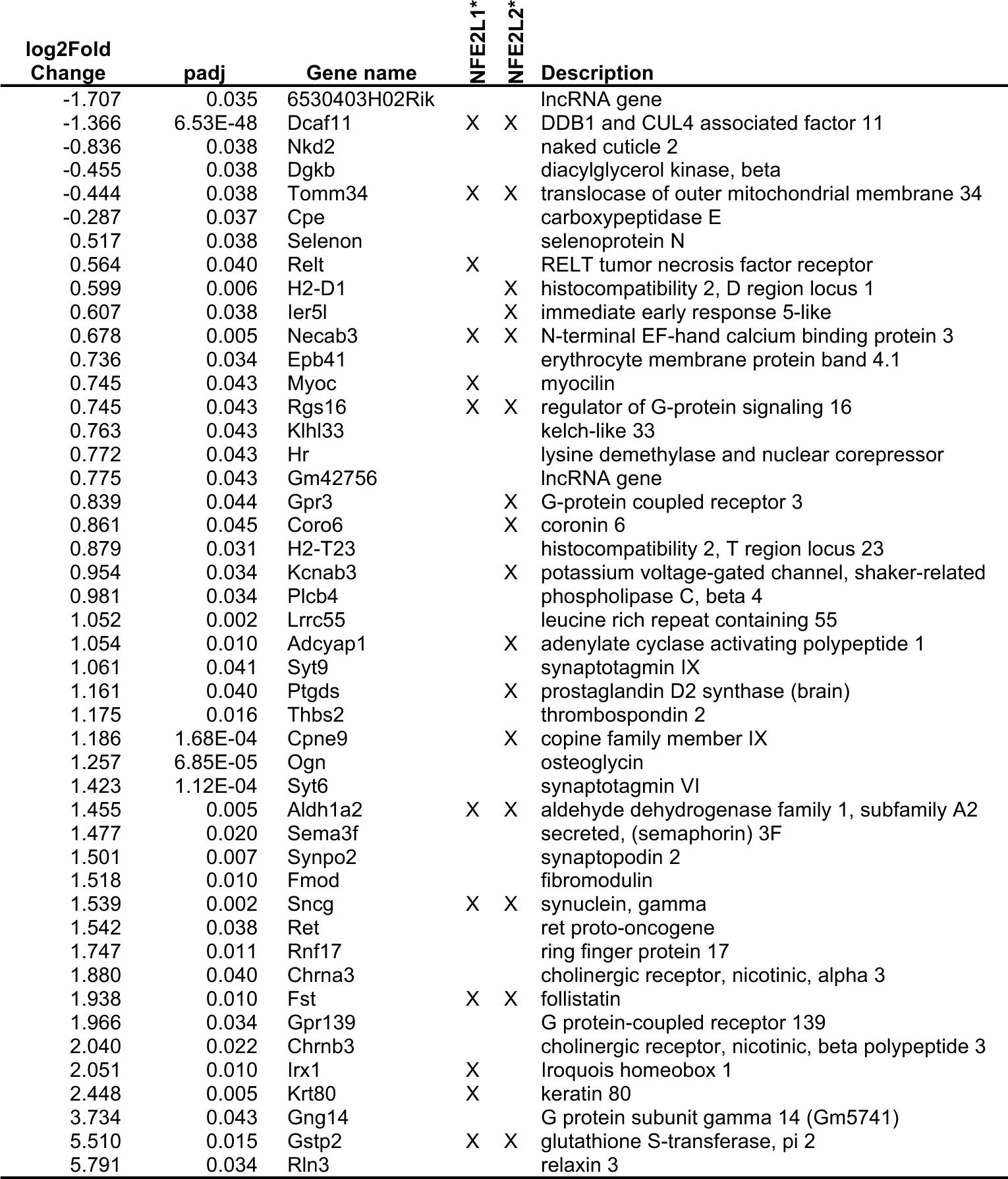
Hippocampus from *Wdr23KO* mice is enriched for NFE2L1/2 target genes. *NFE2L1 and NFE2L2 regulation was identified as previously described [44].

The activity of NRF2/NFE2L2 is regulated by the post-translational degradation of the protein during normal unstressed conditions [40]. We isolated total protein from dissected hippocampus (**Figure 5C-D**) and detected NRF2/NFE2L2 by Western blot analysis. We measured a >150% increase in total NRF2/NFE2L2 protein in *Wdr23KO* (**Figure 5D**), suggesting a steady state stabilization in the absence of stress when WDR23-dependent proteostasis is perturbed [5]. We next isolated and cultured cortical neurons (Figure S5A) and mixed glia (Figure S5B) from WT and *Wdr23KO* mice, which displayed a similar increased stabilization of NRF2/NFE2L2. Taken together, these data confirm the role of WDR23 for the proteostatic control of total NRF2/NFE2L2 in the murine brain, which supports the action for the CUL4-DDB1-WDR23 system working in parallel with CUL3-KEAP1 [41] in the brain as it does in the periphery [4, 5]. Although recent work has demonstrated that female sex hormones do not significantly contribute to sexually dimorphic behavioral responses [42], estrogen has been demonstrated to rapidly activate NRF2/NFE2L2 [43], which is why in this study we look at biochemical and molecular changes in NRF2 activity only in male *Wdr23KO* animals.

### Enhanced capacity for stress resistance in hippocampus of *Wdr23KO* mice

To test whether the increase in established NFE2L2/NRF2-regulated transcripts measured in *Wdr23KO* hippocampus resulted in an increase in steady state abundance of stress adaptation proteins, we quantified the steady state level of several proteins with roles in redox and oxidative stress resistance [45]. We noted an increase in the abundance of several proteins that mediate survival and protection from oxidative stress [46] in the *Wdr23KO* relative to WT; these include the glutathione peroxidase 4 (GPX4, **Figure 6A-B**), glutathione S transferase 4 (GSTA4, **Figure 6A,C**), aldehyde dehydrogenase 2 (ALDH2, **Figure 6A,D**), and the manganese superoxide dismutase 2 [47] (SOD2, **Figure 6E-F**). Intriguingly, we could not detect an increase in all proteins with known sensitivity to NFE2L2/NRF2 activation (Figure S6) suggesting a failure in the sensitivity of detection for these proteins or perhaps specificity to the WDR23-axis of regulation of NFE2L2/NRF2 in the hippocampus specifically, which will be of great future interest.

**Figure 6.**
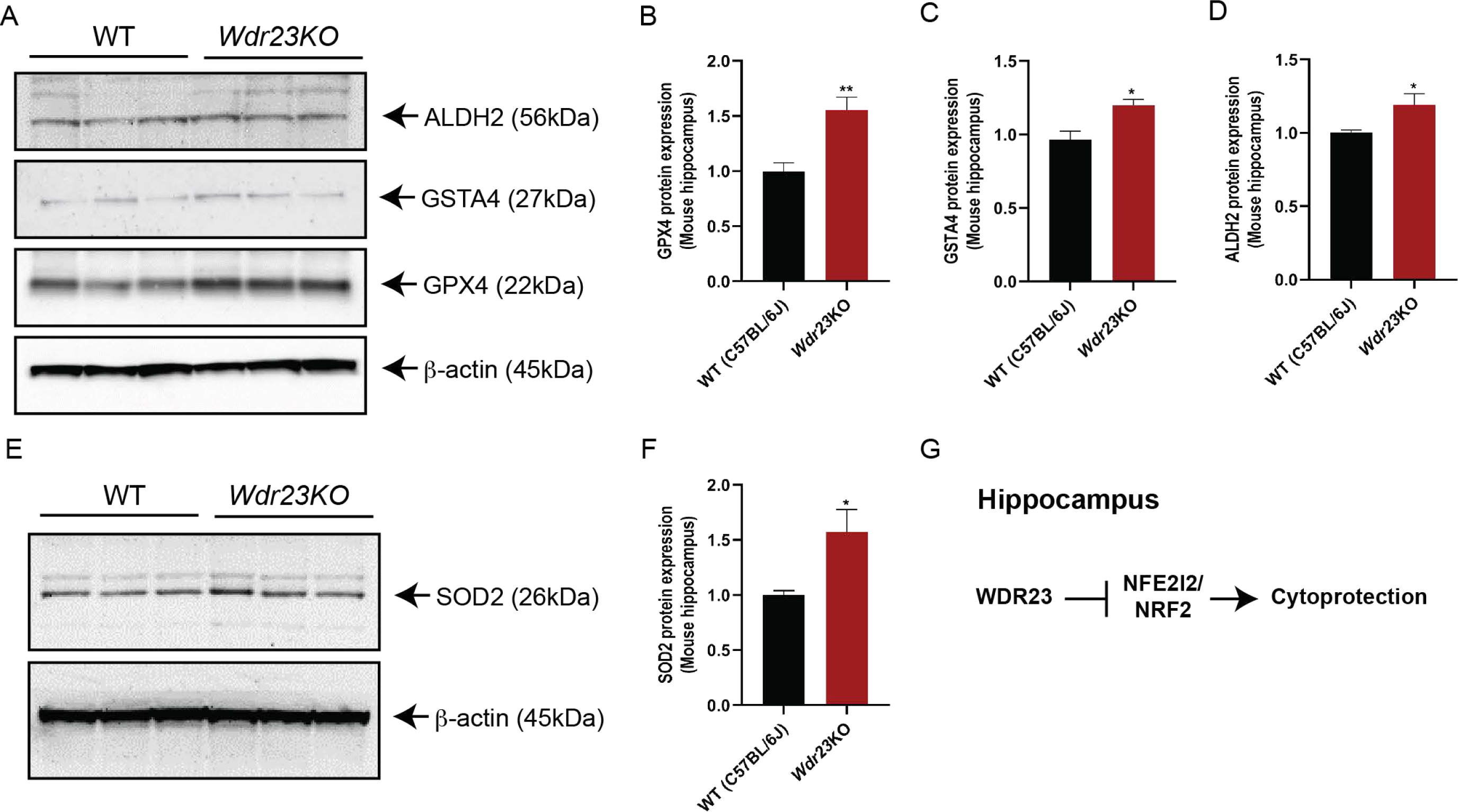
The effects of WDR23 on cellular stress resistance pathways. Loss of *Wdr23* enhanced protein expression of NRF2 target genes. Western blot analysis of GPX4 **(A,B)**, GSTA4 **(A,C)**, ALDH2 **(A,D)** and SOD2 **(E,F)** proteins in hippocampus of male mice. **(G)** The role of WDR23-NRF2 on cytoprotection in hippocampus. *p<.05, **p<.01, ***p<.001, ****p<.0001, compared to age matched WT (C57BL/6J) control. Male mice were fed with chow diet for 10-12 weeks before brain tissue harvesting.

In our study, we uncovered insights into the effects of the loss of *Wdr23* on various behavioral and metabolic parameters. Remarkably, male and female mice often exhibited divergent responses to the loss of *Wdr23*, both in young and older groups. These differences not only highlight the effect of *Wdr23* deficiency in the context of brain aging, but also suggest that sex plays a pivotal role in modulating the impact of *Wdr23* on behavior and metabolism, and underscore the complex interplay between genetic factors, sex, and aging processes, all of which contribute to the intricate landscape of brain aging and its associated phenotypes. Further exploration of these sex-specific differences may unveil valuable insights into the development of targeted therapeutic strategies for gender- and age-specific cognitive decline and neurodegenerative diseases. Collectively, these data support a model where loss of *Wdr23*, stabilizes the steady-state level of NRF/NFE2L2 and results in the increased expression of targets that promote cellular stress resistance (**Figure 6G**), which could influence age-related changes in behavior and be an important new molecular phenotype of successful healthy brain aging.

## ABBREVIATIONS

SAB: spontaneous alternating behavior
NOP: novel object placement
NOR: novel object recognition
ROS: reactive oxygen species
GST: glutathione-s-transferase
SOD: superoxide dismutase
NFE2L2: NFE2 like bZIP transcription factor 2

## SUPPLEMENTAL FIGURE LEGENDS

**Figure S1.**
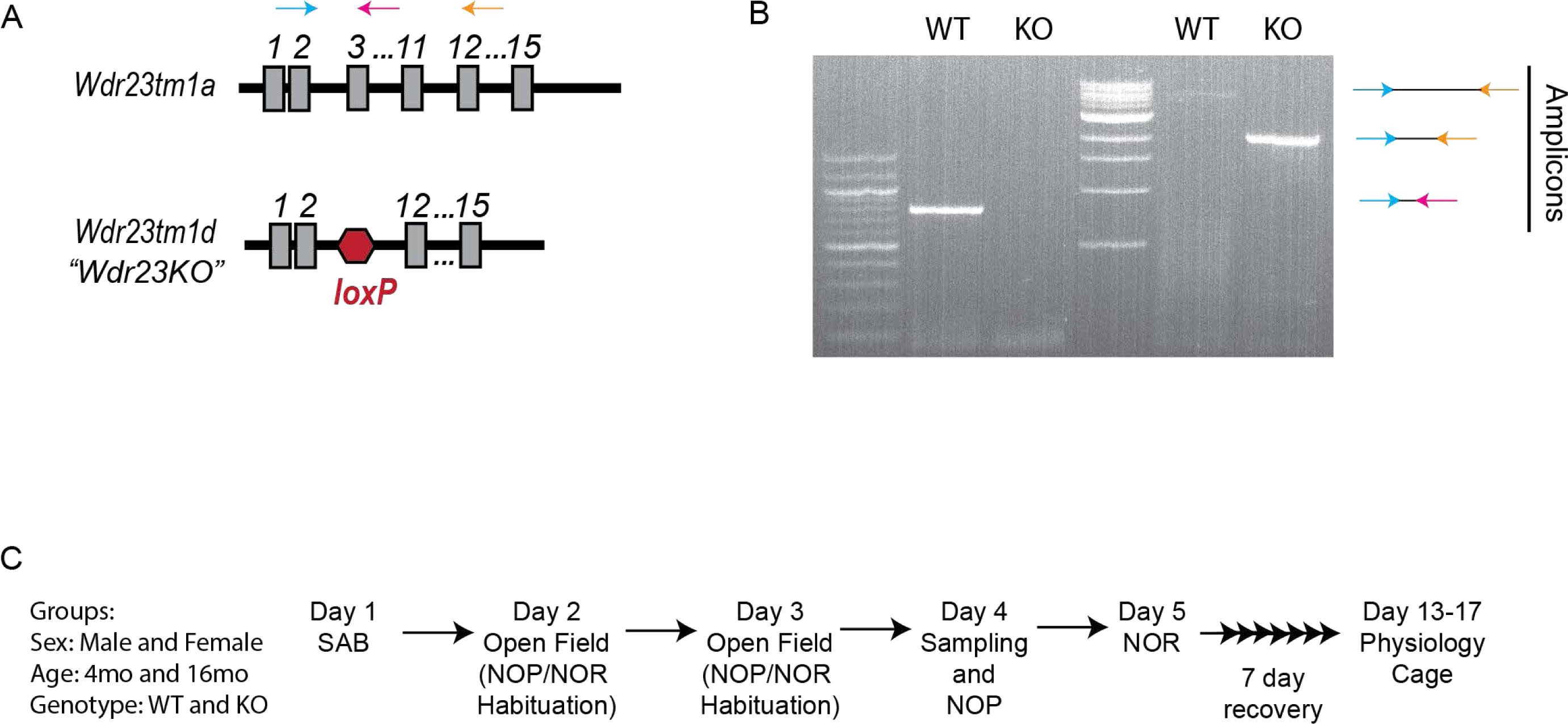
*Wdr23KO* model. **(A)** Cartoon of genetic *Wdr23KO.* **(B)**, Genotyping of WT and *Wdr23KO* samples. **(C)** Experimental design for behavior assessments.

**Figure S2.**
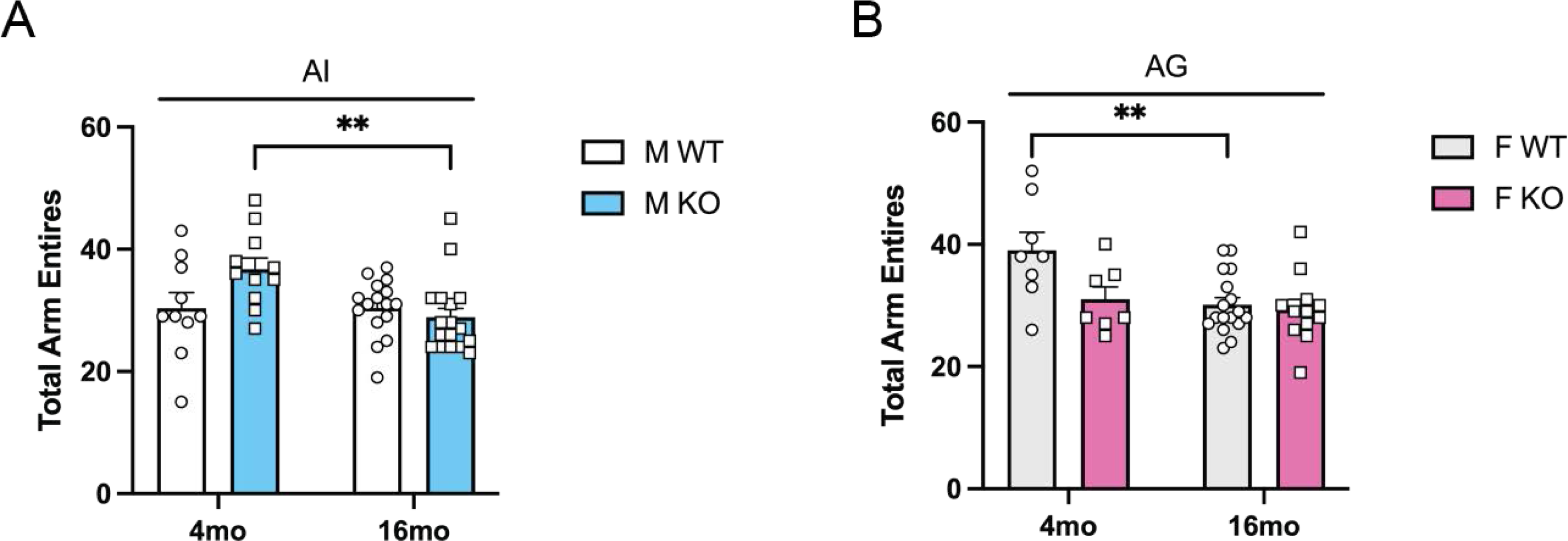
SAB metrics. Total arm entries in the Y maze in **(A)** male and **(B)** female mice. Data are presented as mean (+SEM) values. n=7-17/group. Statistically significant main effects are denoted by A (age), G (genotype) and I (age and genotype interaction). ** denotes p < 0.01.

**Figure S3.**
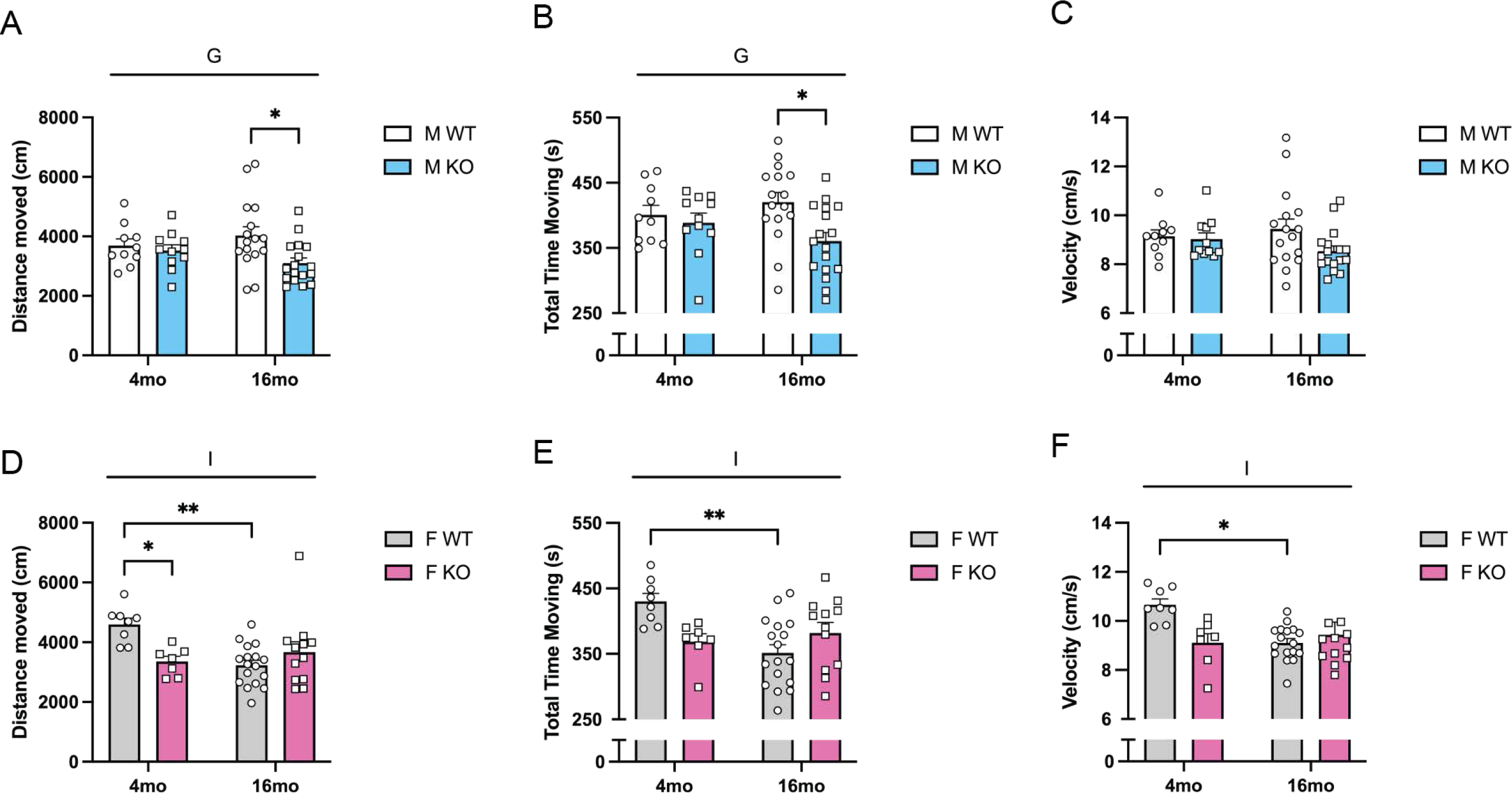
Open field test metrics. **(A, D)** Total distance moved, **(B, E)** the amount of time remained mobile, **(C, F)** the average velocity in the open field test. Data are presented as mean (+SEM) values. n=7-17/group. Statistically significant main effects are denoted by G (genotype) and I (age and genotype interaction). ** denotes p < 0.01.

**Figure S4.**
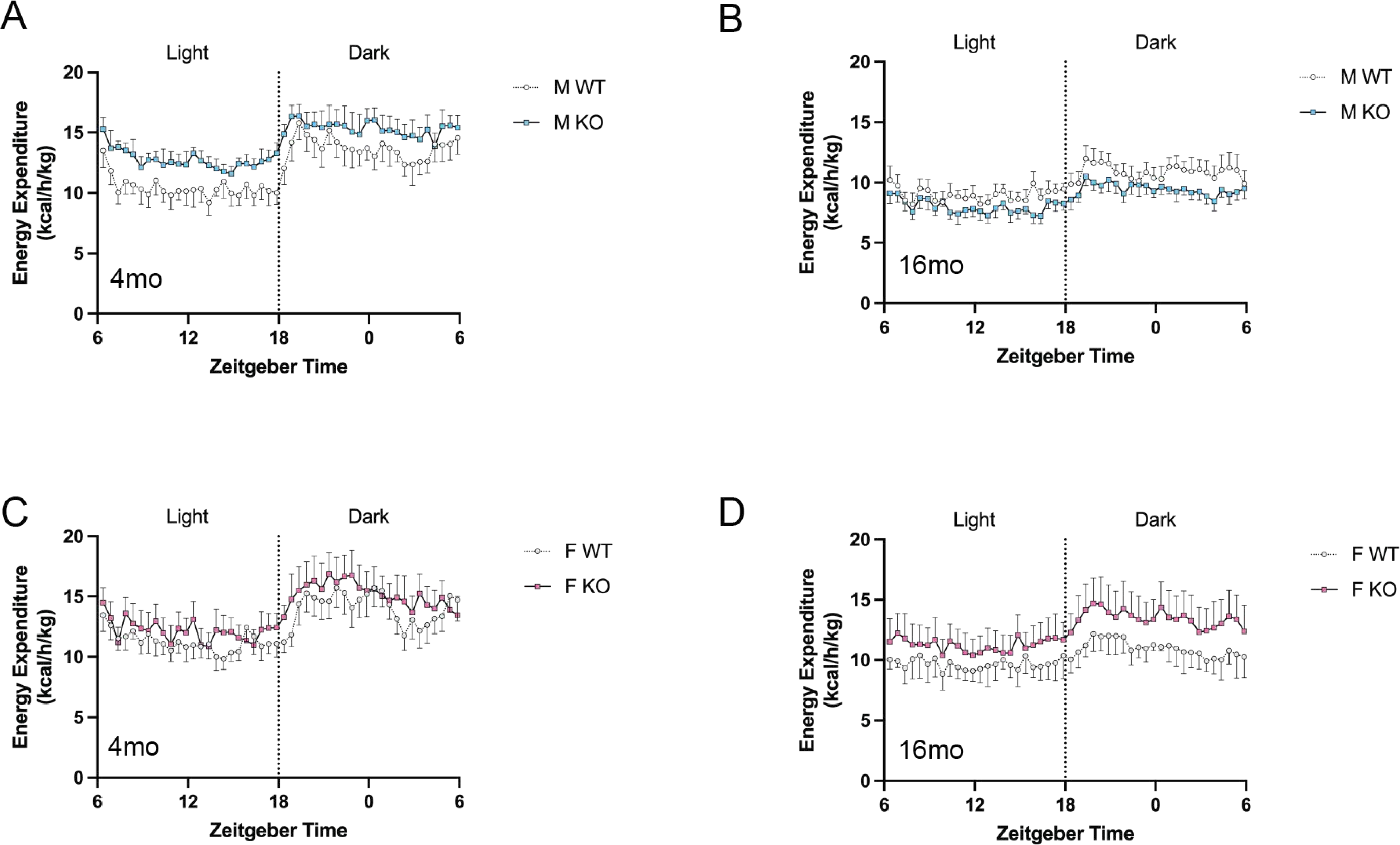
Energy expenditure. Energy expenditure **(EE)** was normalized with body weights. Averaged EE at different time points in **(A)** 4-month-old male, **(B)** 16-month-old male, **(C)** 4-month-old female and **(D)** 16-month-old female mice. Data are presented as mean (+SEM) values. n=4-5/group.

**Figure S5.**
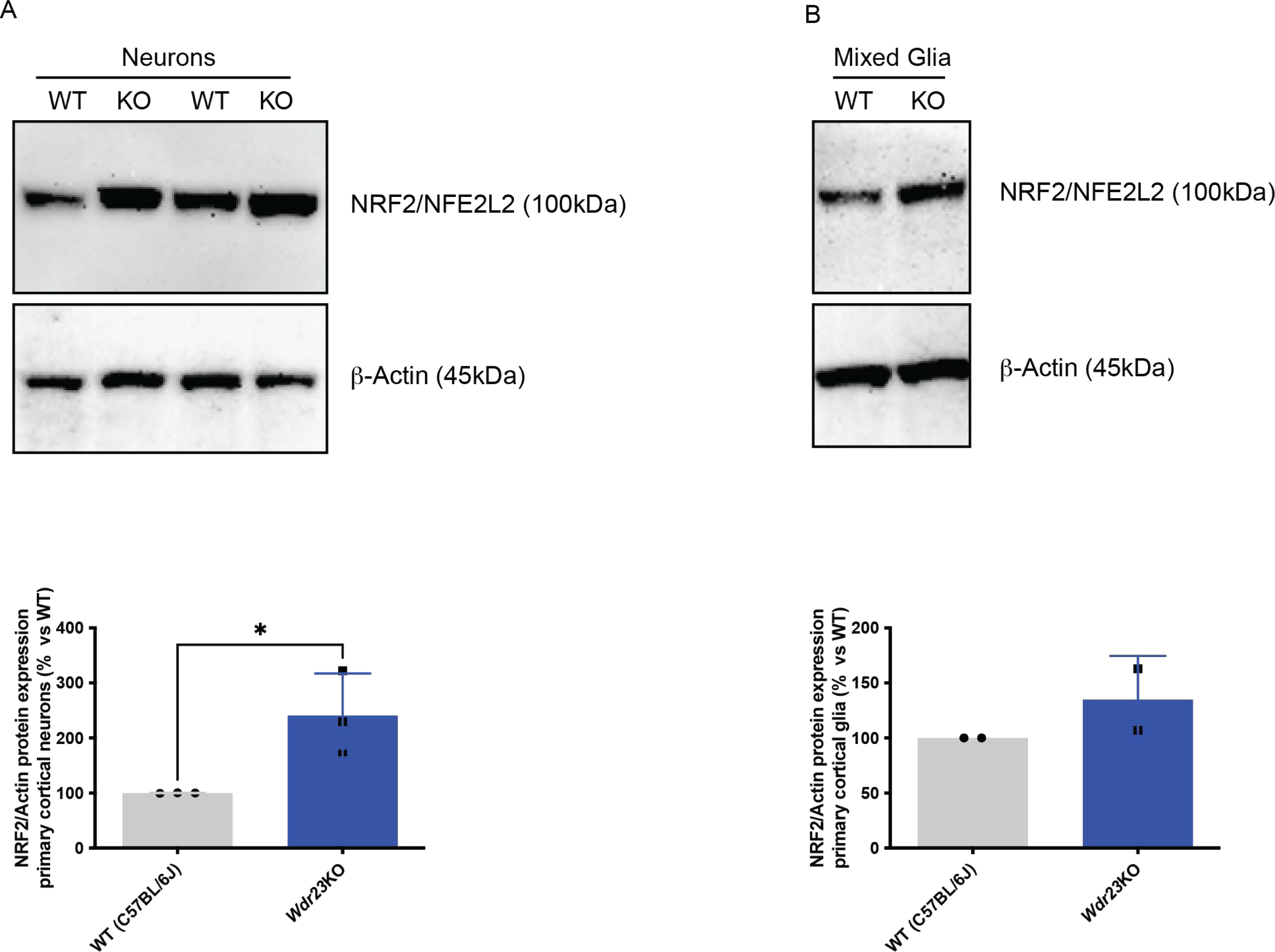
Genetic loss of *Wdr23* increases cortical NFE2L2/NRF2 protein expression. Western blot qualitative analysis of NFE2L2/NRF2 protein in primary cortical neuron culture **(A)**, and primary cortical mixed glial culture **(B)** indicates increased steady-state levels in *Wdr23KO* mouse compared to age matched WT (C57BL/6J) control). n=4-6/group.

**Figure S6.**
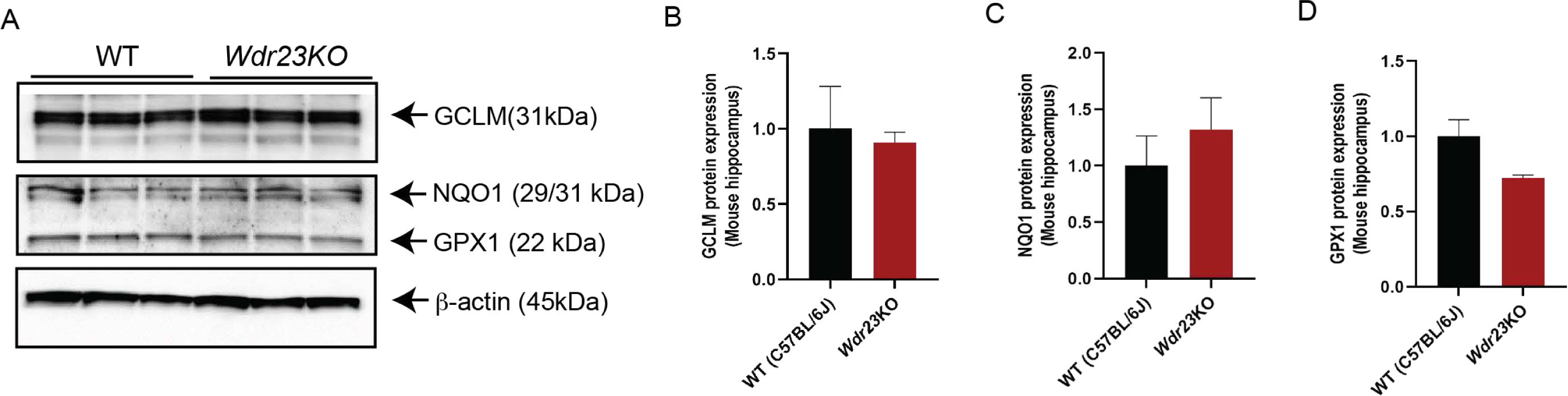
The effects of loss of WDR23 on protein expression of NRF2 target genes. Western blot analysis of GCLM **(AB)**, NQO1 **(AC)**, and GPX1 **(AD)** proteins in hippocampus of male mice. *p<.05, **p<.01, ***p<.001, ****p<.0001, compared to age matched WT (C57BL/6J) control. Male mice were fed with chow diet for 10-12 weeks before brain tissue harvesting. n=3/group

## ACKNOWLEDGEMENTS

We thank M. Donoghue, M. Lynn and S. Ledgerwood for technical assistance. We thank C. Ramos and Dr. N. Stuhr for critical reading of the manuscript and Dr. C. Finch for antibodies recognizing NFE2L2/NRF2 targets. This work was funded by the NIH RF1AG063947 to SPC and a Glenn Foundation for Medical Research Postdoctoral Fellowship in Aging Research from the American Federation for Aging Research to CD.

## Author contributions

Conceptualization: SPC; Methodology: JL, CD, RI, and SPC; Investigation: JL, CD, RI, and SPC; Visualization: JL, CD, and SPC; Supervision: SPC; Writing (original draft): JL and SPC; Writing (reviewing & editing): JL, CD, and SPC.

## Data and materials availability

All data are available in the main text.

